# Rarefaction, alpha diversity, and statistics

**DOI:** 10.1101/231878

**Authors:** Amy Willis

## Abstract

Understanding the drivers of microbial diversity is a fundamental question in microbial ecology. Extensive literature discusses different methods for describing microbial diversity and documenting its effects on ecosystem function. However, it is widely believed that diversity depends on the number of reads that are sequenced. I discuss a statistical perspective on diversity, framing the diversity of an environment as an unknown parameter, and discussing the bias and variance of plug-in and rarefied estimates. I argue that by failing to account for both bias and variance, we invalidate analysis of alpha diversity. I describe the state of the statistical literature for addressing these problems, and suggest that measurement error modeling can address issues with variance, but bias corrections need to be utilized as well. I encourage microbial ecologists to avoid motivating their investigations with alpha diversity analyses that do not use valid statistical methodology.

## 1. Introduction

An alpha diversity metric is a one-dimensional summary of an ecological community. Because many perturbations to a community affect alpha diversity metrics, summarizing and comparing ecologies via alpha diversity is a ubiquitous approach to analyzing community surveys. In microbial ecology, analyzing the alpha diversity of amplicon sequencing data is a common first approach to assessing differences between environments.

Unfortunately, determining how to meaningfully estimate and compare alpha diversity is not trivial. To illustrate, consider the following example where the alpha diversity metric of interest is taxonomic richness (the total number of different taxa present in the environment). Suppose I conduct an experiment in which I take a sample from Environment A and count the number of different microbial taxa present in my sample. I then take a sample from Environment B, count the number of different taxa in that sample, and compare it to the number of taxa in Environment A. I will generally observe higher numbers of different taxa in the sample with more microbial reads, regardless of any true biological differences between the environments. The library sizes dominate the biology in determining the result of the experiment.

Rarefaction is a method that adjusts for differences in library sizes across samples to aid the comparison of diversity. First proposed by Sanders (1968), rarefaction involves selecting a specified number of samples that is equal to or less than the number of samples in the smallest sample, and then randomly discarding reads from larger samples until the number of remaining samples is equal to this threshold. Based on these subsamples of equal size, diversity metrics can be calculated that can contrast ecosystems “fairly,” independent of differences in sample sizes (Weiss et al. 2017).

Unfortunately, rarefaction is neither justifiable nor necessary, a view framed statistically by McMurdie & Holmes (2014) in the context of comparison of relative abundances. In this article I discuss why unequal sample sizes appear to cause special problems in the analysis of diversity in microbial ecology. I introduce a statistical perspective on the estimation of diversity, and argue that ecologists' view of diversity indices is causing fundamental issues in comparing samples. Without advocating for any particular model of microbial sampling, I suggest a new formulation for comparing microbial diversity, one which accounts for uncertainty in estimating diversity metrics. However, since estimates for alpha diversity metrics are heavily biased when taxa are unobserved, comparing alpha diversity using either raw or rarefied data should not be undertaken. I describe statistical methodology for alpha diversity analysis that adjusts for missing taxa, which should be used in place of existing common approaches to diversity analysis in microbial ecology.

## 2. Measurement error and variance in microbiome studies

Imagine that we had complete knowledge of every microbe in existence, including identity, abundance and location. To compare microbial diversity, we would define specific environments (e.g., the distal gut of women aged 35 living in the contiguous U.S.) and compare diversity metrics across different ecological gradients (e.g., with or without irritable bowel syndrome diagnoses). Diversity could be compared exactly, because we would know entire microbial populations with perfect precision.

Unfortunately, we do not have knowledge of every microbe. We take samples from environments, and investigate the microbial community present in the sample. We use our findings about the sample to draw inferences about the environment that we are truly interested in. The samples that we take are not of particular interest except that they are reflective of the environment from which they were sampled. As we sample more and more of the environment using larger samples, we get closer to understanding the true and total microbial community of interest. This means that as we increase sampling, our calculation of any diversity metric (e.g., richness (Fisher et al. 1943), Shannon index (Shannon 1948) and Simpson index (Simpson 1949)) approaches the value of that diversity metric as calculated using the entire population.

Observing small samples from a large population is not an experimental setup unique to microbial ecology: it is almost universal in statistics. The set-up where an estimate of a population quantity converges to the correct value as more samples are obtained is also well understood in statistics. The unique property of microbiome experiments and alpha diversity analysis is that each sample does not faithfully represent the entire microbial community under study. We have ignored and unadjusted error in using our samples as proxies for the entire community.

To illustrate this distinction, I contrast microbial diversity experiments with a classical, non-microbial experimental set-up. Suppose we are interested in modeling the CO_2_ flux of soil treated with different amendments. We would measure the flux of equally sized soil sites treated with the different amendments, performing biological replicates using multiple sites for each amendment. To assess if the amendments affect the flux, we would fit a regression-type model (such as ANOVA) to flux with amendment as an explanatory variable. Implicitly, this model acknowledges that we can assess the flux with high precision; that is, the margin for error for determining flux is negligible.

Now suppose we knew that our flux-measuring machine consistently underestimated flux by exactly 5 units. We would adjust for the measurement error by adding 5 units to each measurement before comparing them. But what happens when we have random measurement error? If the measurement error on the machine was random (e.g., with 0 mean and variance of 1 unit for all amendments), this would not affect any particular amendment. However, detecting a difference between the effects of amendment on flux would be more challenging statistically: we would require more samples to detect a true difference compared to the case without measurement error. To account for the additional experimental noise, we would use a model that would account for measurement error in assessing differences between amendments. If the variance in the measurement error was 1 unit for amendment A but 5 units for amendment B, we would similarly adjust with a measurement error model.

To decide if measurement error must be accounted for when observations are made in an experiment, consider the effect of repeating the observational process on the same experimental unit. In the flux experiment, this would involve measuring the flux of the same soil sites again using the same experimental conditions. Without measurement error in the observations, we would consistently observe the same flux measurement, while if we had random measurement error, we would observe slightly different flux measurements. Because biological replicates in microbiome experiments yield different numbers of reads, different community compositions, and different levels of diversity, we have measurement error in microbial experiments. We generally do not account for this measurement error in microbial ecology studies.

## 3. Bias in estimating and comparing alpha diversity

While measurement error in microbiome studies affects all analyses of microbiome data, alpha diversity is particularly affected because commonly used estimates of alpha diversity are heavily biased compared to other estimation problems in microbial ecology (such as estimating relative abundances). Some tools to address problems with bias in alpha diversity exist in the statistical literature. However, there are two commonly held beliefs about alpha diversity that are preventing the uptake of statistically-motivated methodologies. The first belief is that we are calculating alpha diversity using the best possible estimates of alpha diversity metrics. The second belief is that our alpha diversity calculations should be treated as precisely observed quantities, or known exactly without any measurement error.

To clarify this discussion, I will focus on taxonomic richness (the simplest case), and later generalize the argument to other alpha diversity metrics. Consider the setting in Figure 1a, where we are investigating 2 different environments, and Environment A’s richness (call it *C_A_*) is higher than Environment B’s richness (*C_B_*). Suppose we have two biological replicates of samples from each environment: *n*_*A*1_ and *n*_*A*2_ reads from Environment A, *n*_*B*1_ and *n*_*B*2_ reads from Environment B, and *n*_*A*1_ < *n*_*B*1_ < *n*_*A*2_ < *n*_*B*2_. Let *C_ij_* be the observed richness of environment *i* on replicate *j*. As may commonly occur in practice, *C*_*A*1_ < *C*_*A*2_ < *C*_*B*1_ < *C*_*B*2_.

**Fig 1.**
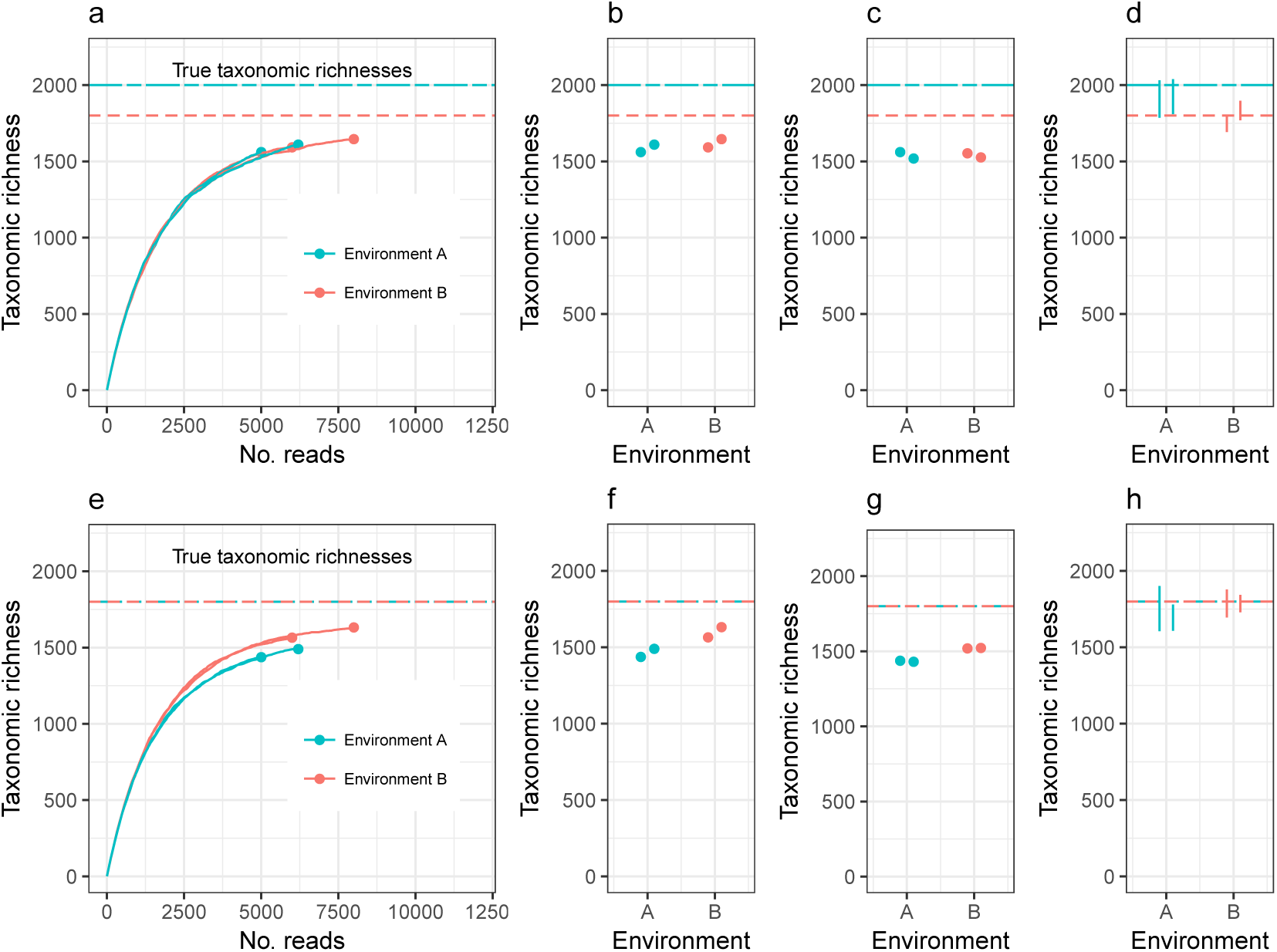
Observed taxonomic richness increases with number of reads (a, e). Comparing raw taxonomic richness can therefore often lead to incorrect conclusions about total diversity (b, f). Rarefying samples to the same number of reads can also lead to incorrect conclusions (c, g). Adjusting for unobserved taxa and accounting for uncertainty in the estimate correctly detects both true (d) and false (h) differences in diversity. While this figure displays taxonomic richness, the same issues apply to other alpha diversity indices.

There are currently two commonly used methods for comparing diversity. The first method, Figure 1b, is to use the estimates *c*_*A*1_, *c*_*A*2_, *c*_*B*1_, and *c*_*B*2_, and perform modeling and hypothesis testing (such as ANOVA or a linear regression) as if both the bias and variance of these estimates were zero (see, for example, Mäkipää et al. (2017)). In the setting of Figure 1a, this leads to the erroneous conclusion that Environment A has lower richness than Environment B. The second method is to generate a normalized, or rarefied sample by randomly discarding reads from all samples until each sample has *n*_*A*1_ reads (the number of reads in the smallest sample), Figure 1c. The resulting rarefied richness levels are then *c*_*A*1_, *c*′_*A*1_, *c*′_*B*1_, and *c*′_*B*2_. These estimates are then used for modeling and hypothesis testing (see, for example, Arora et al. (2017)). This leads to the conclusion that Environment A and Environment B do not have significantly different richnesses. In this case, the estimates of total diversity are far below the actual total diversities of each ecosystem (there is substantial bias in the estimates), prohibiting comparison of diversity across different experiments. Furthermore, not all information collected from the samples was used in making the comparison.

Here I propose and advocate a third strategy: adjust the sample diversity of each ecosystem by adding to it an estimate of the number of unobserved species, estimate the variance in the total diversity estimate, and compare the diversities relative to these errors (Figure 1d). This option has the advantages of leveraging all observed reads, comparing estimates of the actual parameter of interest (total diversity), and accounting for experimental noise. In the case where the environments have equal diversity (Figures 1 e-h), this approach correctly detects equal diversity, even when the abundance structures differ. A Shiny application that compares Type 1 error rates (the probability of incorrectly concluding different levels of total diversity) and Type 2 error rates (the probability of incorrectly concluding equal levels of total diversity) for measurement error modeling approach compared to modeling rarefied or raw alpha diversity is available as Supplementary Materials.

Modeling parameters observed with estimation error is not a new suggestion: this approach is from the field of statistical *meta-analysis*, where the results of multiple studies estimating the same effect size is compared (Demidenko 2004, Willis et al. 2016). In meta-analyses, larger studies need to be given more weight in determining the overall effect size, and this is incorporated into a metaanalysis via the smaller standard errors on the effect size estimates. Similarly, when comparing the response of different treatment groups in clinical trials, the number of subjects in each treatment group is accounted for in a comparison of the overall treatment effect. Adjusting for sample size when comparing different groups of observations without discarding data is widely prevalent in the sciences, and discarding data to adjust for unequal sample sizes is the exception. The strategy outlined here for modeling total diversity after adjusting for missing species adjusts for both bias and variance, thus accounting for library size differences and incomplete microbial surveys.

While the example discussed here is richness, this approach to estimating and comparing alpha diversity using a bias correction (incorporating unobserved taxa) and a variance adjustment (measurement error model) could apply to any alpha diversity metric. However, richness estimation has a well-studied statistical literature, and richness estimators that are adapted to microbiome data exist (Bunge et al. 2014). The same is not true for other alpha diversity metrics. For example, the Chao-Bunge (Chao & Bunge 2002) and breakaway (Willis & Bunge 2015) estimators of species richness provide variance estimates, account for unobserved taxa, and are not overly sensitive to the singleton count (the number of species observed once). In contrast, the coverage adjusted entropy estimator of the Shannon index (Chao & Shen 2003) provides variance estimates and accounts for unobserved taxa, but is extremely sensitive to the singleton count, which is often difficult to determine in microbiome studies. Similarly, the minimum variance unbiased estimate of the Simpson index (Zhang & Zhou 2010) does not account for unobserved taxa. While alpha diversity estimation for microbiomes is an active area of research in statistics (Arbel et al. 2016, Zhang & Grabchak 2016), there remain many features of microbial ecosystems (such as spatial organization of microbes) that are not yet incorporated into statistical methodology for alpha diversity estimation. Despite this, alpha diversity estimates that account for unobserved taxa and provide variance estimates are vastly preferable to both plug-in and rarefied estimates, which do not account for unobserved taxa nor provide variance estimates.

## 4. A call to avoid invalid analyses

Plug-in estimates of many alpha diversity indices (including richness and Shannon diversity) are negatively biased for the environment’s alpha diversity parameter, that is, they underestimate the true alpha diversity when the environment is sampled inexhaustively (Figure 1). Attempting to address this problem using rarefaction (rather than correcting bias with statistically-motivated estimates) actually induces more bias. This is sometimes justified by claiming that rarefied estimates are equally biased. However, this is not generally true, because environments can be identical with respect to one alpha diversity metric, but the different abundance structures will induce different biases when rarefied (Figure 1e shows two environments with different abundance structures but equal richness; rarefying gives the false impression of unequal richness). In this way, both sample diversity and rarefied diversity are not meaningful estimates of a property of the environment, but are driven by an artifact of the experiment (library size). In order to draw meaningful conclusions about the entire microbial community, it is necessary to adjust for inexhaustive sampling using statistically-motivated parameter estimates for alpha diversity. In order to draw meaningful conclusions regarding comparisons of microbial communities, it is necessary to use measurement error models to adjust for the uncertainty in the estimation of alpha diversity.

It has recently been argued that studying microbial diversity without context is distracting us from gaining insight into ecological mechanisms (Shade 2016). To this criticism, I add the argument that misapplying statistical tools is undermining many analyses of alpha diversity. I encourage microbial ecologists to use estimates of alpha diversity that account for unobserved species, and to use the variance of the estimates in measurement error models to compare diversity across ecosystems.

## Acknowledgements

This article is based on course notes presented by the author at the Marine Biological Laboratory at the STAMPS course in 2013, 2014, 2015, 2016 and 2017. The author is grateful to the MBL, the STAMPS course directors, and the STAMPS participants for countless discussions on this topic. The author also thanks Thea Whitman for many thoughtful suggestions on the manuscript.

## Supplementary Materials

Software implementing measurement error error modeling, along with alpha diversity estimates that correct for bias due to unobserved taxa, is available via the R package breakaway. A Shiny application that compares the Type 1 and Type 2 error rates of measurement error models to the error rates for standard approaches to alpha diversity analysis is available as Supplementary Materials.

